# Pseudogene-mediated DNA demethylation leads to oncogene activation

**DOI:** 10.1101/2020.07.15.205542

**Authors:** Junsu Kwon, Yanjing V. Liu, Chong Gao, Mahmoud A. Bassal, Adrianna I. Jones, Junyu Yang, Zhiyuan Chen, Li Ying, Henry Yang, Leilei Chen, Annalisa Di Ruscio, Yvonne Tay, Li Chai, Daniel G. Tenen

## Abstract

Despite being one of the leading causes of cancer-related deaths, there is an unmet clinical need for hepatocellular carcinoma (HCC) patients. The lack of effective treatment is, at least in part, due to our lack of understanding of the molecular pathogenesis of this disease. Oncofetal protein SALL4 is re-activated in patients with aggressive HCC along with other solid tumors and hematologic malignancies. This study identifies a previously unrecognized mechanism of SALL4 reactivation which is mediated by pseudogene-induced demethylation. Using a locus-specific demethylating technology, we identified the critical CpG region for SALL4 expression. We showed that SALL4 pseudogene 5 hypomethylates this region through interaction with DNMT1, resulting in SALL4 upregulation. Intriguingly, pseudogene 5 is significantly upregulated in a hepatitis B virus (HBV) model prior to SALL4 induction, and both are increased in HBV-HCC patients. Our results suggest that pseudogene-mediated demethylation represents a unique mechanism of oncogene activation in cancer.

**Significance:** Our study provides a mechanistic link between HBV infection, activation of the oncogene SALL4, and HCC. We reveal a previously undescribed capability of a pseudogene to epigenetically activate an oncogene by demethylation in a locus-specific manner.

## Introduction

Hepatocellular carcinoma (HCC) is one of leading causes of cancer-related deaths globally, with more than 700,000 new cases and 600,000 estimated HCC deaths each year. Hepatitis B virus infection is one of the main causes of HCC, particularly in Asia. While surgery, liver transplantation, or radiological intervention may be a viable option for early stage disease, prognosis for advanced stage HCC remains bleak, with most patients eventually dying within 20 months after diagnosis. Sorafenib, an oral multikinase inhibitor, is the one of the few approved agents for patients with advanced HCC (1,2). However, the effectiveness of Sorafenib for advanced HCC is debatable (2). There is an unmet clinical need for the development of more effective therapies for the treatment of HCC. The lack of effective treatment options for HCC is at least in part due to our lack of understanding the pathogenesis of this disease. Identifying novel pathway(s) that are responsible for HCC could be translated into targeted therapy and improve the outcomes of these patients.

SALL4 is a potent stem cell factor for self-renewal and pluripotency of embryonic stem cells (ESCs) (3,4). During development, SALL4 expression diminishes gradually and is eventually silenced in most normal tissues. Strikingly, high SALL4 expression levels have been observed in many malignancies such as liver cancer, acute myeloid leukemia, breast cancer and lung cancer (5-8). Re-expression of SALL4 in cancers is associated with a more aggressive cancer phenotype, drug resistance and reduced patient survival (5,6,9-11). Of note, HCC patients with detectable SALL4 expression have enriched hepatic progenitor-like gene signatures and poorer prognoses (10). Furthermore, targeting SALL4 in HCC cell lines by knocking down or using inhibitory peptides resulted in cellular death (12), suggesting that SALL4 plays a crucial role in hepatocarcinogenesis and that targeting SALL4 may provide an innovative therapeutic approach for this disease. However, mechanistically, how SALL4 is re-activated in HCC is still unclear, although it has been reported that aberrant methylation could be a contributing factor (13,14). By defining the mechanism of SALL4 reactivation in HCC, we can better treat HCC.

DNA methylation is a frequently studied mechanism of epigenetic regulation in humans that is mediated by DNA methyltransferases (DNMT); of which, DNMT1 has a structural binding preference (15). Research by multiple groups including ours has demonstrated that non-coding RNAs (ncRNAs) such as ecCEBPα, Dali, Dum, and Dacor1 can interact with DNMT1 to inhibit its methylation activity. These ncRNAs thus indirectly alter local methylation states in different cancers, acting as key tissue-specific epigenetic regulators of gene expression (15-18). It was also reported that the exon 1-intron 1 region of the *SALL4* gene locus is hypermethylated in non SALL4-expressing K562 leukemic cells. Reprogramming of these cells resulted in demethylation of this region and a subsequent increase in SALL4 expression (14). Recently, a report described demethylation of specific CpG sites downstream of the SALL4 transcriptional start site (TSS) in hepatitis B (HBV)- related HCC which could contribute to SALL4 re-activation in HCC (13). However, it is still unclear how HBV infection could initiate the demethylation and reactivation of specific oncogenes.

Pseudogenes are a class of non-coding RNAs (ncRNAs) once regarded as insignificant “junk” DNA relics due to their lack of coding potential. However, studies have demonstrated that pseudogene transcripts can regulate gene expression of oncogenes and tumor-suppressors by acting as antisense transcripts, processed small interfering RNAs (siRNAs) and competing endogenous RNAs (ceRNAs) (19-21). Recent pseudogene expression analysis in over 2,800 patient samples showed strong concordance between pseudogene expression and tumor subtypes, as well as patient prognoses, highlighting the clinical importance of pseudogenes (22). Our group focused on characterizing regulatory functions of SALL4 pseudogenes. SALL4, a well-studied oncogene with high expression levels in several hematological malignancies and solid tumors, has eight pseudogenes of different lengths varying from 500 nucleotides to 6,000 nucleotides, and yet there have been no studies investigating SALL4 pseudogenes (5-8).

As many pseudogenes are actively transcribed in cells, we postulated that they could interact with RNA-binding proteins such as DNMT1 via highly homologous RNA motifs and exert regulatory functions. We therefore tested the hypothesis as to whether pseudogenes are involved in DNA methylation as DNMT1-interacting lncRNAs in an HBV-positive HCC model.

## Results

### SALL4 expression is negatively correlated with methylation of the 5’ UTR - exon 1 - intron 1 region

We hypothesized that the degree of methylation in the SALL4 locus is associated with SALL4 expression, therefore, we examined HBV-positive HCC patients. Using a publicly available dataset (23), the overall transcript levels and methylation status of SALL4 were analysed using probes covered the entire SALL4 gene locus. A significant negative correlation between SALL4 expression and methylation was observed in primary tumours at the Probe 1 (Fig. 1A and 1B) which was only observed in the 5’UTR – exon 1 region. Sites located either proximal or distal (Probe 2) to the 5’UTR-exon 1 locus showed poor to no correlation with SALL4 expression (Fig. 1C).

**Figure 1:**
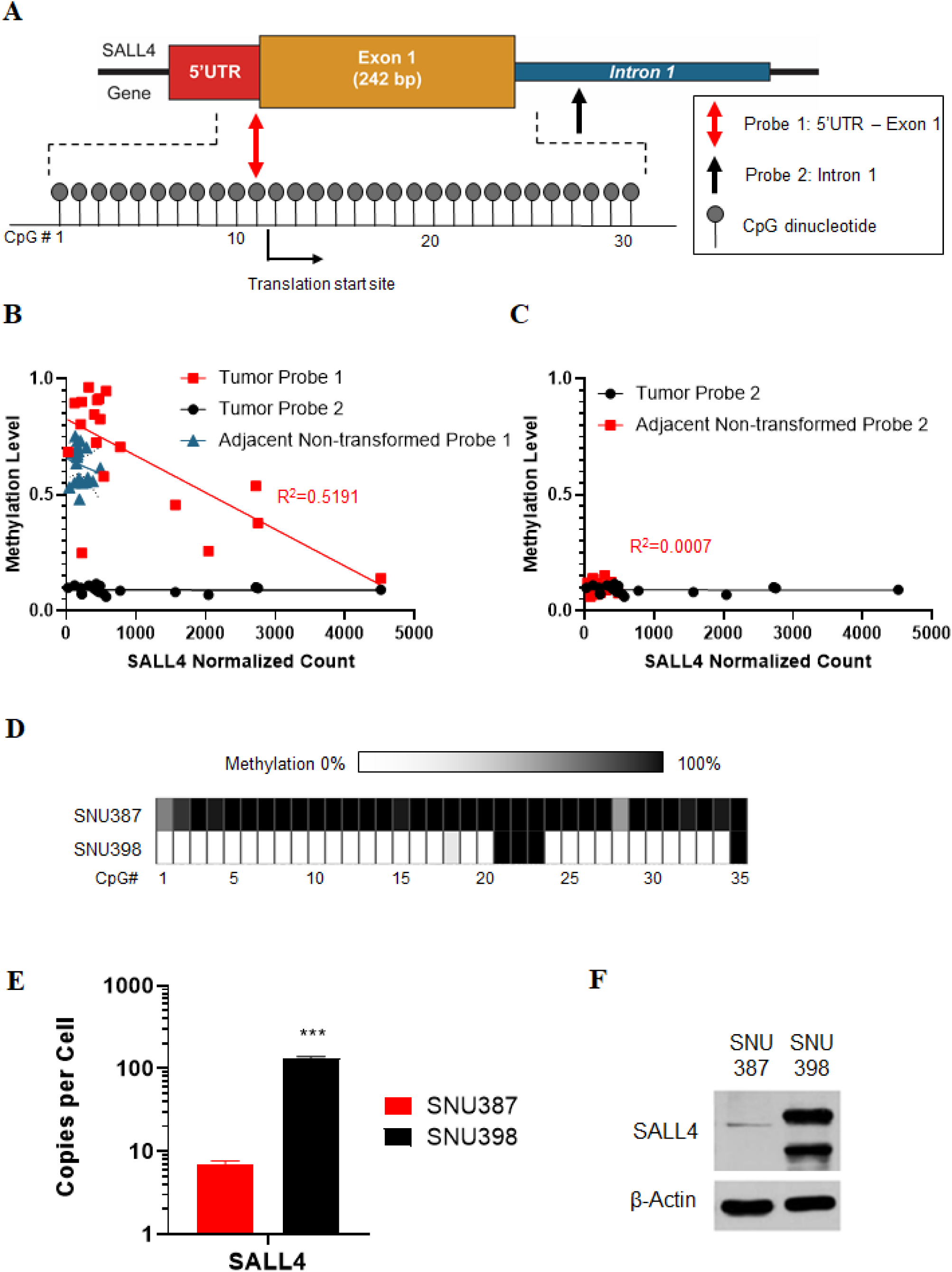
SALL4 expression is negatively correlated with methylation of the 5’UTR-exon 1-intron 1 region. (**A**) Schematic representation of the methylation probes. The numbers refer to each CpG dinucleotide. The probe in the 5’UTR-exon 1 junction, “Probe 1”, assesses the methylation status of the CpG dinucleotide #11 and the intronic probe, “Probe 2” assesses the CpG dinucleotide #68 in Supplementary Figure 1. (**B and C**) SALL4 expression and methylation correlation analysis in 19 HBV+ patients. Compared to paired adjacent non- transformed liver tissue, there is a negative correlation between SALL4 expression and Probe 1 methylation, which is not observed using Probe 2. (**D**) Bisulfite sequence of the 5’UTR-exon 1 intron 1 region in wildtype SNU398 and SNU387 HCC cell lines. White color represents hypomethylation while black represents hypermethylation of the individual CpG dinucleotide. Degree of methylation was determined as a proportion of methylated cytosine residue at a position out of 10 clones. Only CpG dinucleotides 1 to 35 are represented as the sequencing efficiency was poor for dinucleotides 36 to 39. (**E**) Absolute quantification of SALL4 mRNA expression in wildtype SNU398 and SNU387. β-actin was used as a positive control for the assay. cDNA for β-actin quantification was diluted 10 times and back- calculated accordingly later. The levels of β-actin were comparable between SNU398 and SNU387 at about 400 to 600 copies of transcripts per cell. However, SNU398 cells expressed more than 150 copies of SALL4 mRNAs while SNU387 expressed less than 10 copies on average. (**F**) SALL4 protein levels in wildtype SNU398 and SNU387. β-actin was used as a loading control for immunoblotting.

As the methylome and transcriptome of cell lines could be different from those of primary tumours, we further investigated the negative correlation between SALL4 methylation and expression among HCC cell lines. The 5’UTR-exon 1-intron 1 region was first inspected and found to have over 30 CpG dinucleotides (Supplementary S1). Bisulfite sequencing in the HCC cell lines SNU398 and SNU387 revealed distinct and unique methylation profiles for the two cell lines (Fig. 1D). Within the profiled region, SNU387 showed a near universal, methylated profile in stark contrast to SNU398 which showed a completely demethylated profile, with the exception of 4 CpG dinucleotides. The result was consistent with previous reports observing that methylation of the SALL4 5’UTR-exon 1-intron 1 region is differentially methylated in K562-induced pluripotency reprogrammed cells and HBV-related HCC patients. (13,14). To investigate the relationship between the observed methylation profiles and gene expression, we examined SALL4 expression in both SNU398 and SNU387 (Fig. 1E and 1F). The level of SALL4 transcription was substantive as more than 100 copies of SALL4 mRNAs per cell were detected in SNU398, while SNU387 cells only expressed about 10 copies per cell. It was also evident that SALL4 was expressed at much higher magnitude in SNU398, in which the SALL4 loci was hypomethylated. Taken together, both the cell line and patient data suggest that SALL4 methylation and expression are negatively correlated. We therefore confirmed that DNA methylation could be a potential regulatory mechanism for SALL4 expression in HCC.

### CRISPR-DiR demethylates and activates SALL4

To further investigate the correlation between methylation of the SALL4 5’ UTR- exon 1- intron 1 region and SALL4 expression, the CRISPR-DNMT1-interacting RNA (CRISPR-DiR) technique was utilized to induce locus-specific demethylation by blocking DNMT1 activity in SNU387 cells. Briefly, the single guide RNA (sgRNA) was constructed to contain a SALL4 exon 1 targeting sequence, two RNA loops of ecCEBPα with validated DNMT1 inhibitory function (15), and the dCAs9-interacting domain (Fig. 2A). Numerous sgRNAs targeting the SALL4 5’ UTR- exon 1- intron 1 locus were designed, with the four most efficient candidates shortlisted (sgSALL4_1 - sgSALL4_4) via *in vitro* sgRNA selection (Supplementary Fig. S2A). Transduced cells were also validated to express the dCas9-mCherry through FACS (Supplementary Fig. S2B).

**Figure 2.**
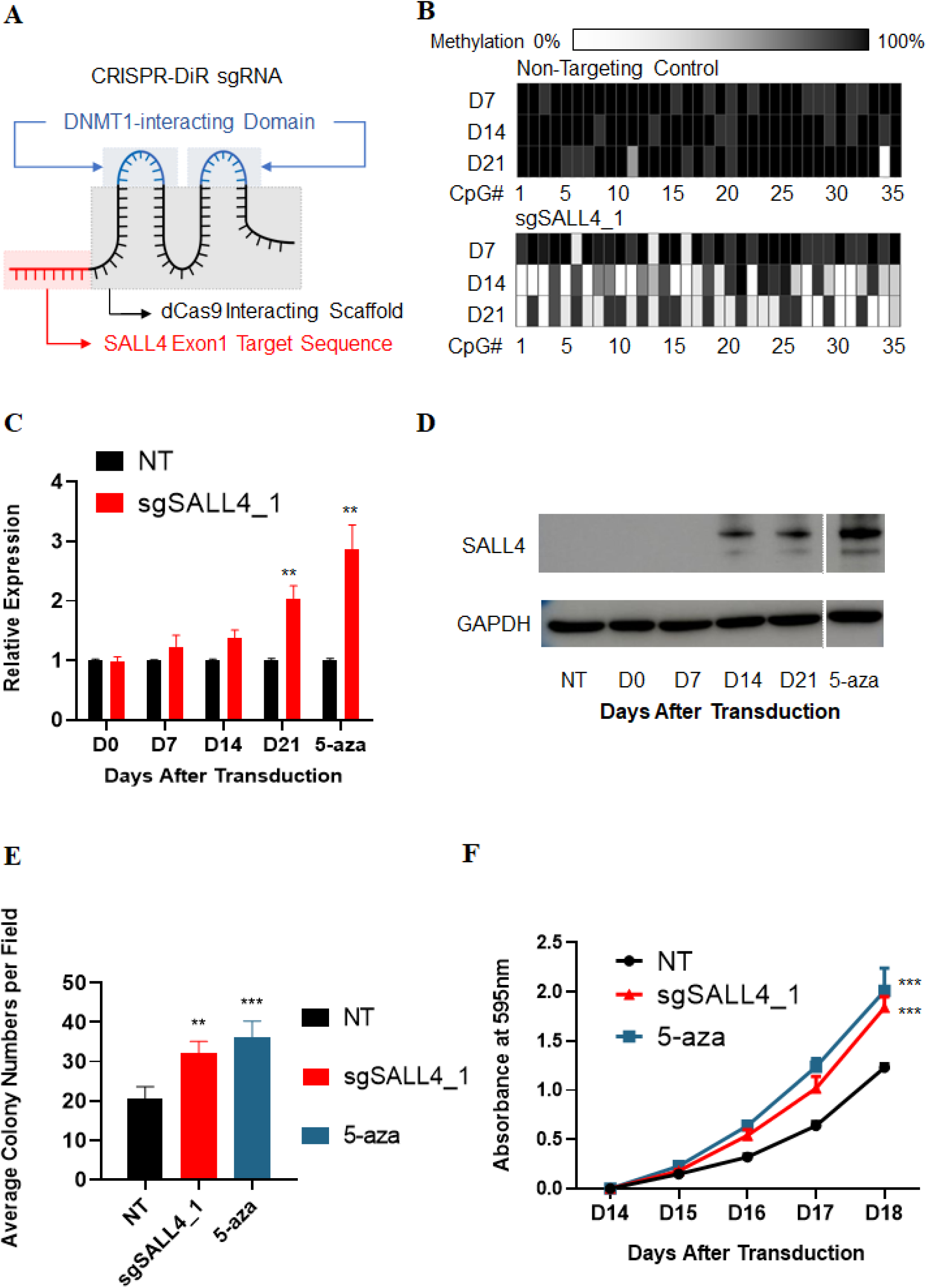
CRISPR-DiR demethylates and activates SALL4. (**A**) single-guide RNA design for the CRISPR-DiR. The red region targets and interacts with the SALL4 5’UTR. The black region interacts with dCas9. The blue regions are the two segments of ecCEBPα that interact with DNMT1 (15). (**B**) Bisulfite sequence of CRISPR-DiR transduced SNU387 cells. The data represents the methylation profile 14 days after CRISPR-DiR for SALL4. The numbers indicate each of the CpG dinucleotides in the 5’UTR-exon 1 intron 1 junction. The white color represents hypomethylation while the black hypermethylation of the individual CpG dinucleotides. Only CpG dinucleotides 1 to 35 are represented as the sequencing efficiency was poor for dinucleotides 36 to 39. (**C** & **D**) SALL4 transcript and protein levels after CRISPR-DiR in SNU387. The western blot image is cropped as there were multiple lanes in between “D21” and “5-aza”. However, they are from the same blot and exposed for the same duration. (**E)** Soft agar growth assay for CRISPR-DiR in SNU387. (**F**) Growth curve assay for CRISPR-DiR for SALL4 in SNU387. 5-aza-2-deoxycytidine(decitabine) was used as a positive control. NT denotes non-targeting negative control. Mean ± SD, n ≥ 3, *P < 0.05; **P < 0.01; ***P < 0.001.

Methylation of the SALL4 5’ UTR- exon 1- intron 1 CpG island was monitored in SNU387 cells with four independent CRISPR-DiR inductions, one for each shortlisted sgRNA. Of these inductions, sgSALL4_1 was the most potent sgRNA tested. Upon transduction of SNU387 cells with sgSALL4_1, significant demethylation changes were observed after 14 days, which continued for over 7 additional days (Fig. 2B). Conversely, no change in methylation was observed in non-targeting, negative control transduced cells. To examine potential off-target effects of CRISPR-DiR, we concurrently monitored methylation of a region in SALL4 exon 4, and confirmed demethylation was localized to only the targeted 5’ UTR - exon 1 - intron 1 CpG island (Supplementary Fig. S2C).

Following CRISPR-DiR targeted demethylation of the SALL4 5’ UTR-exon 1-intron 1 CpG island, both SALL4 transcript and protein levels increased as predicted (Fig. 2C and 2D). The magnitude of SALL4 upregulation observed in these cells was comparable to that of treatment with 5-aza-2’-deoxycytidine, a global demethylating agent. Furthermore, SALL4 expression was not significantly altered when the cells were transduced with other less efficient sgRNAs (Supplementary Fig. S2D). As SALL4 overexpression promotes cancer cell growth, we performed growth assays on sgSALL4-transduced SNU387 cells and observed increased anchorage-independent and -dependent growth compared to negative control (Fig. 2E and F), suggesting that targeted demethylation of the SALL4 locus leads to upregulated expression of SALL4, with concomitant enhanced cellular growth.

### SALL4P5 demethylates and activates SALL4, and associates with DNMT1

There are 8 SALL4 pseudogenes, and since none are located on the same chromosome as SALL4, it is unlikely that they will be transcribed as siRNAs or antisense transcripts to deregulate SALL4. However, the identified pseudogenes do share high sequence homology with the paralogous coding SALL4 gene. It is therefore possible that the SALL4 pseudogenes could bind to other proteins with either matching DNA/RNA motifs or with comparable secondary and tertiary structures owing to their high sequence homology. As previously reported, ecCEBPα, a ncRNA that overlaps with and thus has regions of identity with its paralog, CEBPα, can interact with DNMT1 and affect CEBPα gene expression. We therefore postulated that SALL4 pseudogenes, which are highly homologous to SALL4, could potentially mediate SALL4 demethylation.

Each SALL4 pseudogene was transiently overexpressed in SNU387 cells, and the methylation profile of the 5’ UTR-exon 1-intron 1 CpG island was assessed (Fig. 3A). Only SALL4 pseudogene 5 (SALL4P5) overexpression resulted in a demethylation pattern comparable to that seen using CRISPR-DiR. Consistently, SALL4P5 knock-down in hypomethylated SNU398 cells led to increased methylation of the locus as predicted (Fig. 3B).

**Figure 3.**
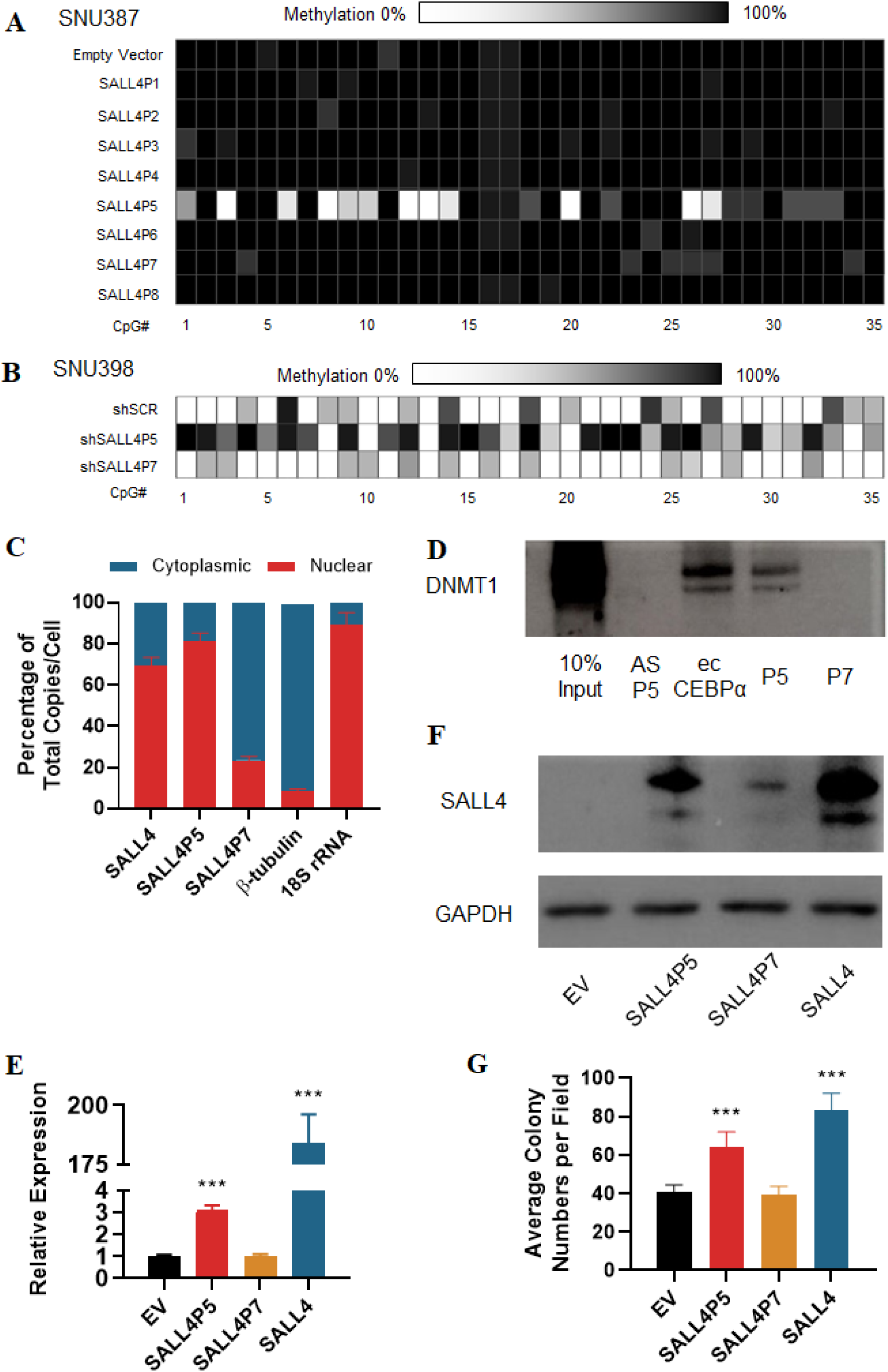
SALL4P5 demethylates and activates SALL4, and associates with DNMT1. (**A**) Bisulfite sequencing after transient overexpression of individual SALL4 pseudogenes in SNU387. The methylation status of CpG dinucleotides in SALL4 5’UTR-exon 1 intron is shown. (**B**) Bisulfite sequencing after SALL4P5 knockdown in SNU398. ShScr denotes scrambled shRNA. shSALL4P7 was used as a negative control. (**C**) Transcript localization in SNU398. Cells were fractionated into nuclear and cytoplasmic fractions and transcript expression was quantified. β−tubulin was used as a cytoplasmic fraction control, 18S rRNA as the nuclear fraction control. (**D**) Biotin-labelled pull-down of DNMT1 in SNU398. Full length SALL4P5 was used as a bait to pull down complexes and DNMT1 presence was probed using immunoblotting. “as P5” denotes the negative control, antisense-SALL4P5 and “ecCEBPα” denotes the positive control. Full length SALL4P7 was used as a pseudogene negative control as well. (**E** and **F**) SALL4 transcript and protein expression after pseudogene overexpression in SNU387. (**G**) Soft agar growth assay for pseudogene overexpression in SNU387. Mean ± SD, n ≥ 3, *P < 0.05; **P < 0.01; ***P < 0.001.

Additional evidence to suggest direct SALL4P5-DNMT1 interaction can be seen by their matched cellular localization. Cellular localization of pseudogene transcripts is a critical factor in determining their function, as they must be localized in the same cellular compartment as their binding partners to exert specific biological functions. It is known that DNMT1 facilitates methylation exclusively in the nucleus. Critically, SALL4P5 is also primarily localized to the nucleus. This contrasts with SALL4P7, which has a predominant cytoplasmic localization in SNU398 (Fig. 3C). Therefore, although SALL4P7 shares sequence homology with SALL4P5 and SALL4, its primary localization in cytoplasm could, in part, account for its inability to demethylate the SALL4 locus.

As DNMT1 is known as a maintenance DNA methylator, we therefore investigated whether our observed SALL4P5 demethylation phenotype is due to a SALL4P5-DNMT1 interaction. As there are no known interacting SALL4P5-DNMT1 binding regions or pockets, we performed an unbiased biotinylated pull-down assay using full-length SALL4P5. First, to validate the efficacy of the pull-down, DNMT1 protein could successfully pull-down ecCEPBα (Fig. 3D). Similarly, SALL4P5 was able to successfully pull-down DNMT1, whereas SALL4P7 and the antisense negative control did not.

Having shown an association between SALL4 exon 1 - intron 1 demethylation and SALL4 expression up-regulation, we next investigated the effect of transiently overexpressing SALL4P5 on SALL4 levels. SALL4P5 overexpression significantly upregulated SALL4 transcript (Fig. 3E) and protein levels, the latter of which was more striking and equivalent to the overexpression of SALL4 itself (Fig. 3F). Interestingly, SALL4P7 overexpression also increased SALL4 protein levels. Although SALL4P7 may not play a role in SALL4 demethylation, it could still contribute to gene regulation as homologous pseudogenes could also function as ceRNAs (21) by sequestering bioavailable microRNAs that target and repress SALL4. Consistent with the phenotype of elevated SALL4 levels, SALL4P5 overexpression also significantly increased colony formation of SNU387 cells (Fig. 3G). The data suggest that SALL4P5 could have oncogenic effects as it can directly upregulate SALL4 expression and cell growth.

### SALL4P5 is upregulated in HCC patients and during hepatitis B induction

The aforementioned results demonstrate that SALL4P5 upregulation can reactivate SALL4 expression in cell lines. We next sought to validate these findings in primary patient samples, and measured SALL4P5 expression in HCC patients, who frequently have elevated levels of SALL4 (10). Twenty HCC patients with paired non-disease samples were screened from a Hong Kong cohort. Among these 20 patients, 19 of them were HBV positive and 7 had increased SALL4 levels of over 1.5-fold (Fig. 4A). Interestingly, within these 7 patients, only SALL4P5 expression was concomitantly upregulated, while SALL4P7 showed little to no change. For patients, such as patient #1, with no SALL4 level change, SALL4P5 expression was also unaltered.

**Figure 4.**
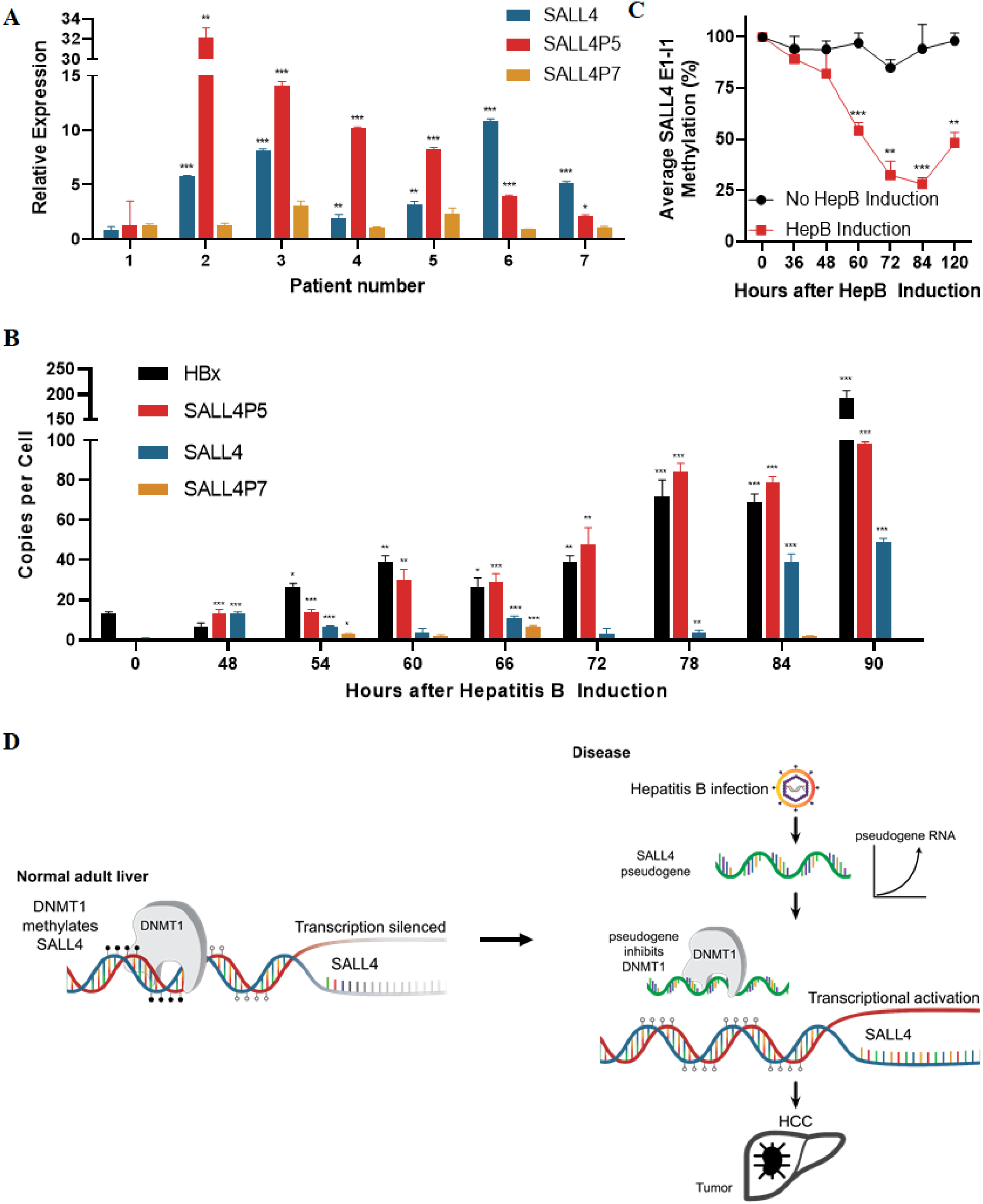
SALL4P5 is upregulated in HCC patients and during hepatitis B induction. (**A**) Relative SALL4 transcript expression in paired HCC patient samples. All expression data are normalized against adjacent non-transformed tissues. Patient #1 was used as a negative control with unaltered SALL4 and SALL4P5 levels. The levels of SALL4, SALL4P5, and SALL4P7 were assessed for the other six patients, patients #2 to 7, as they had more than 1.5-fold elevation in SALL4 levels compared to adjacent non-transformed tissued. (**B**) Absolute quantification (RNA copies per cell) of Hepatitis B antigen X, HBx, and SALL4 transcripts during hepatitis B induction in HepAD38B. Transcript levels of HBx and SALL4 transcripts were monitored every 6-12 hours post HBV induction (**C**) Average methylation in SALL4 5’UTR-exon 1 intron 1 across the 35 CpG dinucleotides in HepAD38B. Blue denotes SALL4 methylation profile without HBV induction, while red denotes SALL4 methylation profile after induction. Mean ± SD, n ≥ 3. *P < 0.05; **P < 0.01; ***P < 0.001.

HBV infection is the single most common risk factor of HCC, as more than 50% of patients contract hepatitis B prior to HCC (24). We therefore sought to investigate whether SALL4P5-mediated demethylation and subsequent reactivation of SALL4 during hepatitis B infection could drive oncogenesis. The HepAD38B model was used, as it allows controlled induction of hepatitis B virus production using the tet-off system (13). Using digital droplet PCR (ddPCR), we validated that the HepAD38B cells produced the major hepatitis B viral transcripts such as core, surface, hepatitis B antigen X (HBx) and polymerase transcripts (Supplementary Fig. 3A). Upon hepatitis B induction, SALL4 and SALL4P5 transcript levels also increased (Fig. 4B, Supplementary Fig. S3B and S3C). Interestingly, the expression level for HBx increased first, SALL4P5 expression then followed at 54 hours, and lastly SALL4 expression at 84 hours in step-wise manner. When performing bisulfite sequencing of these critical time points, it was found that the average methylation across the CpG island in the 5’UTR - exon 1 – intron 1 junction decreased upon hepatitis B induction, suggesting that the infection-induced upregulation of SALL4P5 demethylates and reactivates SALL4. (Fig. 4C).

## Discussion

In this study, we provided a mechanistic link between HBV infection, activation of the oncogene SALL4, and HCC development. We, for the first time to our knowledge, showed that hepatitis B viral infection could lead to pseudogene upregulation, which in turn could epigenetically regulate oncogenes and drive tumorigenesis. We also demonstrated for the first time that a pseudogene can associate with a DNA methyltransferase to inhibit its function and subsequently influence expression of its coding paralog, oncogenic SALL4. We described a strong negative correlation between SALL4 expression and exon 1-locus specific methylation in HCC patients as well as in cell lines. By utilizing a novel CRISPR-DiR technology, we validated the importance of exon 1 methylation to SALL4 expression and cell growth. Interfering with DNMT1 activity at a specific CpG island in this region resulted in the upregulation of SALL4. Furthermore, we identified and characterized SALL4 pseudogene 5 (SALL4P5), which shares high sequence homology with its paralogous coding gene SALL4, as a critical regulator of SALL4 expression and function by interacting with DNMT1 to demethylate and upregulate SALL4 expression. More importantly, we demonstrated that SALL4P5 and SALL4 expression are sequentially upregulated in an HBV induction model as well as positively correlated and upregulated in hepatitis B-infected HCC patients.

This work highlights the previously undescribed capability of a pseudogene to epigenetically regulate an oncogene by interacting with DNMT1 and affecting its methylation in a specific region. It is reported that there are at least 12,000 pseudogenes (25) in the human genome. A recent pan-cancer analysis of pseudogene expression in different cancers demonstrated that pseudogene expression alone can serve as a molecular and prognostic factor for patients with different cancer subtypes, highlighting the functional and clinical importance of pseudogenes (22). There may be other pseudogenes capable of mediating similar DNMT1-interactions and exerting the demethylating function on oncogenes and tumor-suppressors in different cancers. Monitoring expression levels of oncogene- demethylating pseudogenes could enable predicting oncogene activation as well as disease progression to improve patient outcomes.

Understanding the mechanism of pseudogene-mediated demethylation may provide insights into SALL4 re-expression in HCC, which is of therapeutic value owing to SALL4’s prognostic significance for the disease (7). Here we elaborate how a pseudogene could play a crucial role in epigenetic regulation of an important oncogene, thereby suggesting that there could similarly significant and impactful pseudogenes potentially contributing to early-stage gene regulation of tumor-suppressors and oncogenes. In addition, pseudogenes could exert non-canonical functions by interacting with other RNA-binding proteins, with potential wide-ranging implications in gene regulation and function as we have demonstrated here.

Previous studies from Di Ruscio et al., demonstrated that RNAs require distinct secondary structures in order to associate with DNMT1 (15). This preferential interaction through structure could potentially explain why DNMT1 interacts with SALL4P5, but not SALL4P7, even though the two pseudogenes are highly homologous. Another striking aspect of SALL4P5-SALL4 regulation is that unlike ecCEBPα, which resides in the CEBPα locus, the SALL4P5 locus is on chromosome 3, while its paralogous coding gene SALL4 is on chromosome 20. This *trans* regulation implies that it could be the sequence homology and perhaps secondary structure that plays a more critical role than chromosomal location for a ncRNA to work as a DNMT1-interacting RNA. Moreover, as DNMT1 plays a crucial role in *de novo* methylation, the SALL4P5-DNMT1 interaction could contribute to SALL4 reactivation as well as other oncogene activation in cancers.

DNA hypermethylation of CpG islands in gene promoter regions has consistently correlated with inactivation of tumor suppressors in cancers (26). Conversely, demethylation of oncogene promoters leads to increased gene expression (27). Interestingly, it was the methylation profile of the 5’UTR - Exon 1-Intron 1 region of SALL4 that was critical in SALL4 upregulation leading to cell growth. Further investigation will be needed to determine whether this non-canonical methylation site, downstream from the promoter, is only significantly affected in the context of pseudogene-mediated demethylation.

These studies represent one of the first examples of gene locus specific demethylation resulting in the activation of an oncogene. By an innovative strategy, we have identified the specific CpG island in the locus that is required for SALL4 activation and expression. This approach could be extended to other loci to identify the CpG “regulating” modules allowing gene expression and controlled by an RNA-mediated mechanism.

Our studies suggest a model in which hepatitis B viral infection upregulates SALL4P5, followed by SALL4 (Fig. 4D). This novel insight addresses unmet clinical need in HCC as HCC is one of the leading causes of cancer-related deaths globally and chronic hepatitis B virus infection accounts for more than 50% of HCC cases (1,28). Elucidating molecular mechanisms of oncogene activation during hepatitis B virus infection could enhance our understanding of the pathogenesis of HCC, and hence, aid in development of robust therapies. Increased SALL4P5 expression is observed in HBV-related HCC patients, making this pseudogene and its related function in oncogene SALL4 activation relevant to developing novel therapeutics in HBV-related HCC.

## Methods

### Cell Culture

All HCC cell lines were obtained from ATCC and grown according to the manufacturer’s instructions in the absence of antibiotics. Human hepatocellular carcinoma cell lines (SNU398, SNU387 and HepAD38B) were maintained in Dulbecco’s Modified Eagle’s medium (DMEM) and Roswell Park Memorial Institute 1640 medium (RPMI) (Life Technologies, Carlsbad, CA) with 10% fetal bovine serum (FBS) (Invitrogen) and 2 mM L- Glutamine (Invitrogen). These cell lines were cultured at 37°C in a humidified incubator with 5% CO2. SNU398 was derived in 1990 from a 42-year-old, Asian male hepatocellular carcinoma patient. SNU387 was derived in 1990 from a 41-year-old, Asian female hepatocellular carcinoma patient. HepAD38B cell line was derived from a 15-year-old, male hepatoblastoma patient.

### RNA extraction and gene expression analysis

Total RNA was extracted from cells using the Trizol® reagent (Invitrogen) and purified using the RNeasy Mini kit from Qiagen. 1 µg of purified RNA was used for cDNA synthesis using the High-Capacity cDNA Reverse Transcription Kit (ThermoFisher Scientific) according to the manufacturer’s instructions. The QuantStudio 5 Real-Time PCR System (Thermo Fisher Scientific) was used to assess the expression levels of the mRNAs, miRNAs, and pseudogenes of interest. GoTaq® qPCR Master Mix (Promega) was used as a SYBR master mix reagent for the qPCR procedures. The qRT-PCR data was analyzed using the QuantStudioTM Design & Analysis Software Version 1.2 (ThermoFisher Scientific) and represented as relative expression(ΔΔCt), normalized against either GAPDH or β-actin. The primer sequences used for the quantitative real-time PCR are provided in Supplementary Table 1.

### Genomic DNA extraction

Genomic DNA was extracted from HCC cell cultures using DNeasy Blood & Tissue kit(Qiagen) for bisulfite-sequencing assay according to the manufacturer’s protocols.

### Bisulfite treatment and sequencing

SALL4 5’UTR Exon 1 3’UTR region methylation status was assessed using bisulfite sequencing. In brief, 1 μg of genomic DNA extracted using the DNeasy Blood & Tissue kit(Qiagen) was bisulfite-converted by using the EZ DNA Methylation kit (Zymo Research). PCR products were gel-purified (Qiagen) from the 1.5% TAE gel and cloned into the pGEM-T Easy Vector System (Promega) for transformation. The cloned vectors were transformed into ECOS 101 DH5α cells and miniprep was performed to extract plasmids. Sequencing results were analysed using BiQ analyser software. Samples with more than 90% conversion rate and 70% sequences identity were analysed. The minimum number of clones for each sequenced condition was 6. Primers used for the bisulfite sequencing of SALL4 5’ UTR - exon 1 - intron 1 are listed in Supplementary Table 2.

### Protein extraction and immunoblotting

Total cell lysates in protein lysis buffer (PLB) (100mM KCl (Ambion), 5mM MgCl2 (Ambion), 25mM EDTA pH 8.0 (Life Technologies), 10mM HEPES (Life Technologies), 0.5% NP-40 (Roche), 20mM DTT(Fermentas), Proteinase inhibitor tablet (Roche)). PLB was added to the cell pellets and incubated for 15 minutes on ice. The lysates were centrifuged for 10 minutes at 15,000 x g at 4 °C. Protein concentrations were measured using the Bradford Protein Assay (Bio-Rad Laboratories) and absorbance was measured at 595 nm on the Tecan Infinite® 2000 PRO plate reader (Tecan, Seestrasse, Switzerland). Equal amounts of protein for each sample were diluted with 4X sample buffer (ThermoFisher Scientific) and heated at 95 °C for 5 minutes. The proteins were resolved by SDS-PAGE 12% self-cast gel and transferred onto polyvinylidene difluoride (PVDF) membrane using the Mini Trans-Blot® Electrophoretic Transfer Cell (Bio-Rad) in transfer buffer (25 mM Tris, 192 mM Glycine, and 20% (v/v) methanol (Fischer Chemical)]. After blocking with 5% milk in Tris-NaCl buffer (TBS) (ThermoFisher Scientific) with 0.1% Tween-20 (Sinopharm Chemical Reagent Co., Ltd) (TBST), membranes were incubated with the appropriate primary and secondary antibodies. The immune-reactive proteins were detected using the protein bands were visualized using SuperSignalTM West Dura Extended Duration Substrate (ThermoFisher Scientific) and visualized on Image Quant LAS 500 machine (GE Healthcare) according to the manufacturer’s instructions. SALL4 (Santa Cruz Biotechnology, EE30, #sc-101147), GAPDH (Cell Signaling Technology, #5174), and DNMT1 (Abcam, #ab87656) antibodies were used for immunoblotting as per manufacturer’s instructions.

### Bacterial transformation

ECOSTM 101 competent cells(DH5α) from Yeastern Biotech Co., Ltd. were used for transformation following the manufacturer’s instructions. 50μl of cells were thawed at room temperature in a water bath. 2μl of pre-chilled DNA was immediately added. The tubes were kept on ice for 5 minutes to increase the transformation efficiency. The cells went through heat shock in a 42°C water bath for 30 seconds. The cells were kept on ice again for 5 minutes and plated on LB plates. The plates were incubated at 37°C for 16 hours.

### Plasmid extraction

Plasmid was extracted from 1.5 ml of bacterial culture with the QIAamp DNA Mini Kit(Qiagen) and purified for sequencing validation (1st BASE). For transfected cell line DNA extraction, plasmid was extracted from the cell pellet that is suspended in 100 μl of PBS.

### Plasmid transfection

SNU398 cells were seeded at a density of 75,000 cells/well in 12-well plates 24 hours before transfection. SNU387 and SNU182 cells were seeded at higher density of 100,000 cells/well in 12-well plates. 500 ng of plasmid was added to 3 µl of P3000 reagent (Life Technologies) in 75 µl of Opti-MEM prior to mixing with 2 µl of Lipofectamine 3000 (Life Technologies). The reagent mixture was incubated at room temperature for 10 minutes before adding to each well.

### Cell viability assay

24 hours post-transfection, cells were trypsinized and split into 5 individual wells of 5 separate 12-well plates. Upon adherence, cells were fixed using 10% neutral buffered formalin solution (Sigma-Aldrich, HT501128-4L) and labelled as day 0. Subsequently, the remaining plates were fixed daily from day 2 to day 5 (excluding day 1) prior to staining with crystal violet (Sigma-Aldrich, C0775-100G) for 3-5 minutes at room temperature. Stained wells were washed three times with Milli-Q water and left to dry. Crystal violet stain was solubilized using 10% acetic acid (Sigma-Aldrich, A6283-2.5L). The plates were left on a shaker at room temperature for at least 20 minutes. The absorbance reading for each well was measured at 595 nm using the Tecan Infinite® 2000 PRO plate reader (Tecan, Seestrasse, Switzerland).

### Soft agar assay

A 0.6% agarose base was prepared by mixing 3.9 ml of 2% Ultrapure agarose (Invitrogen) with 9.1 ml of cell culture medium. 2 ml of the mixture was added to individual wells of 6- well plates. 24 hours post-transfection, cells were trypsinized, counted, and diluted to a concentration of 15,000 cells/well. 450 µl of the 2% agarose was added to 2.55 ml of cells for a final agarose concentration of 0.3%, and 1 ml of the mixture was added to each well containing the solidified 0.6% agarose base. 1 ml of culture media was added to the top agar layer upon solidification and the cells were incubated at 37 °C in a humidified incubator.

Culture medium was changed every two days. Images of the colonies were taken under 4X magnification every 4-5 days for a period of up to 14 days and quantified using ImageJ.

### *In vitro* transcription (IVT) and biotinylated RNA pulldown

The DNA template was first amplified by PCR with primers containing a 5’ T7-tag for in vitro transcription. Antisense SALL4P5 control was also amplified by having the reverse primers carrying the T7-tag. The IVT was performed as per manufacturer’s guidelines. 1 μg of purified PCR product was incubated with the transcription mix which was composed of 10X transcription buffer, 400 mM NTP mix, and 200 U T7 RNA polymerase for 5 hours at 37 °C. 140 μl of RNase-free water and 1000 μl of 100% ethanol were added to the transcription product and incubated for at least 30 minutes at -20 °C. The reaction mixture was centrifuged for 1 hour at 4 °C to precipitate the RNA. The RNA pellet was collected and dissolved in 100 μl of ultra-pure water. The RNA was further purified using RNeasy Mini 250 columns (Qiagen) according to the manufacturer’s instructions. The purified RNA obtained from IVT was labelled with biotin at the 3’end using the PierceTM RNA 3′End Desthiobiotinylation Kit (ThermoFisher Scientific) according to the manufacturer’s protocol. Biotin labelling efficiency of the RNA probes was determined using the Chemiluminescent Nucleic Acid Detection Module Kit (ThermoFisher Scientific) following the manufacturer’s protocol. Biotin labelling efficiency was normalized against the efficiency of antisense transcript to determine amount of initial RNA bait used for the subsequent pulldown experiment. Pulldown using these labelled RNA probes was carried out using the PierceTM Magnetic RNA-Protein Pull-Down Kit (ThermoFisher Scientific) according to the manufacturer’s instructions. Protein lysates eluted from the pulldowns were used for immunoblotting and other downstream analysis. The primer sequences with the T7-tag for the PCR are provided in Supplementary Table 3.

### *In vitro* generation of sgRNA transcripts

Approximately 1.4kb of genomic fragment spanning SALL4 5’ UTR - exon 1 - intron 1 was PCR amplified (Zymo Research) and cloned into the pGEM-T Easy vector. The vector was linearized with BamH1 restriction enzyme (New England Biolabs). SALL4-targeting sgRNA candidates were transcribed with HiScribe™ Quick T7 High Yield RNA Synthesis Kit (New England Biolabs) following manufacturer’s instructions. The sgRNA target sequences within SALL4 locus can be supplied upon request

### *In vitro* cleavage and selection of sgRNA transcripts

In vitro cleavage assay was performed using purified Cas9 nuclease from S. pyogenes (New England Biolabs) in order to select SALL4-specific sgRNA among candidates. The experiment was performed according to the manufacture’s protocol. The sgRNAs were denatured at 95°C for 3 minutes, then Cas9 protein and sgRNAs were incubated for 10 minutes at 25°C to form a complex. Lastly, a linearized SALL4 DNA fragment was added to the mixture and the entire reaction was incubated at 37°C for 1 hour. The reaction mixture was composed of purified Cas9 protein, individual sgRNA, and linearized SALL4 genomic fragment in ratio of 10: 10: 1. 1 ul of Proteinase K was added to each sample after the cleavage reaction, and it was then incubated at room temperature for 10 minutes. The result was analyzed with a 1% agarose gel.

### Lentiviral transduction of sgSALL4_1 and dCas9

Lentiviruses expressing dCas9 or sgRNA were packaged in 293T cells the plasmids psPAX2 and pMD2.G. TransIT-LT1 Transfection Reagent (Mirus) was used for transfection into 293T cells. Virus was collected at 48 hours and 72 hours post-transfection. The collected virus was filtered through 0.45 µm microfilters and stored at -80 °C. Transduction of SNU- 387 cells was performed by mixing virus and 4 μg/mL polybrene (Santa Cruz) together to add to the cells seeded in T75 flasks 24 hours prior to the transduction. 24 hours after the transduction, the medium was replenished with normal RPMI culture medium. Transduction efficiency was determined by GFP (for sgRNA) or mCherry (dCas9) expression by FACS analysis, and the positive cells were sorted by a FACS Aria machine (BD Biosciences).

### 5-aza-2’-deoxycytidine(decitabine) treatment

SNU387 cells were treated with 1.25 uM of 5-aza-2’-deoxycytidine (Sigma-Aldrich) according to the manufacturer’s instructions. Culture medium and drug were refreshed every 24 hours due to the drug being light-sensitive. RNA (for RT–PCR) and genomic DNA (for bisulphite sequencing) were isolated after 5 days of treatment

### Digital droplet PCR

Reactions for the ddPCR were prepared by harvesting 100,000 cells on each day for RNA extraction and cDNA preparation. The reaction mixture was prepared with the 2x ddPCR supermix for probes (Biorad, Cat #186-3026), 10-fold diluted cDNA, nuclease-free water, and forward and reverse primers. Once the reaction mixture was ready, it was loaded into a DG8 cartridge for the QX200 Automated Droplet Generator (Biorad, Cat#186-4003). We then proceeded to the thermal cycling with a Biorad C1000 Touch Thermal Cycler with the following cycle conditions: 95°C for 10minutes, 94°C for 30 seconds (40 cycles), 60°C for 2 minutes (40 cycles), 98°C for 10 minutes and 4°C hold. The reaction plate was loaded into the QX200 Droplet Reader (Biorad, Cat#186-4003) for gene expression analysis.

### SALL4 methylation and expression correlation analysis

Pearson correlations between SALL4 expression and methylation levels were performed for the sites within the 5’ UTR - exon 1 - intron 1 intron 1 region and distant sites in the intron 1 (Fig S2). In order to show that a negative correlation is specific to primary HBV+ HCC patients, adjacent normal samples were used as negative controls. The data used for the correlation was taken from Yang, et al (15), containing 19 pairs of primary HBV+ HCC patients and their matched adjacent normal tissues.

## Materials Availability

All plasmids and mouse lines generated in this study are freely available from the authors upon reasonable request.

## Data and Code Availability

The data used for the correlation was taken from Yang, et al, 2017 containing 19 pairs of primary HBV+ HCC patients and their matched adjacent normal tissues. The authors declare that all other data supporting the findings of this study are available within the paper and its supplementary information files.

## Declaration of interests

The authors declare no competing interests.

This study was supported by Singapore Ministry of Health’s National Medical Research Council; National Institutes of Health; Xiu Research Fund, National Research Foundation Fellowship; National University of Singapore President’s Assistant Professorship; Singapore Ministry of Education under its Research Centres of Excellence initiative; Singapore Ministry of Education’s AcRF Tier 3 grants; National Cancer Institute; Italian Association for Cancer Research (AIRC).

## Author contributions

D.G.T., L.C., and Y.T. initiated the project and provided guidance throughout. D.G.T, L.C., Y.T., C.G., and J.K. designed the experiments. J.K. carried out experiments, analyzed data, prepared figures, and wrote the manuscript. C.G, Y.L, and A.J carried out experiments, prepared figures and edited the manuscript. M.A.B. analyzed the data, prepared figures, and edited the manuscript. H.Y. and L.Y. performed the bioinformatics analysis on SALL4 expression and methylation. A.D.R. and J.Y. reviewed the manuscript. D.G.T, L.C, and Y.T conceived of and supervised the project, designed experiments, and critically reviewed the manuscript.

## Acknowledgements

We thank Sudhakar Jha and Polly Chen for their insightful suggestions. We also thank the Tenen, Tay, and Chai lab members for reviewing the manuscript. D.G.T. is funded by the Singapore Ministry of Health’s National Medical Research Council under its Singapore Translational Research (STaR) Investigator Award, as well as NIH grants 1R35CA197697 and P01HL131477. Y.T. is funded by a Singapore National Research Foundation Fellowship and National University of Singapore President’s Assistant Professorship. This research is supported by the National Research Foundation Singapore and the Singapore Ministry of Education under its Research Centres of Excellence initiative, as well as the RNA Biology Center at the Cancer Science Institute of Singapore, NUS, as part of funding under the Singapore Ministry of Education’s AcRF Tier 3 grants, Grant number MOE2014-T3-1-006.

L.C. is funded by NIH/NHLBI grant P01HL095489. A.D.R. is funded by NCI R00CA188595, the Italian Association for Cancer Research (AIRC) start up grant #2014- 15347, and the Giovanni Armenise-Harvard Foundation.

## Notes

### Competing Interest Statement

The authors have declared no competing interest.

