## Supplementary Figures 1-4, Tables 1-4 for "Pseudogene-mediated DNA demethylation leads to oncogene activation"

**Supplementary Figure Legends**

**Supplementary Figure 1. Schematic representation of CpG islands within the 5’UTR‑exon 1 intron 1 region of SALL4**. The numbers refer to each CpG dinucleotide. The ‘ATG’ codon highlighted in bold denotes the translation start site. Underlined nucleotides refer to the primer sequences used for the subsequent bisulfite sequencing experiments. Red, yellow, and blue shades denote SALL4 5’UTR, exon 1, and intron 1, respectively

**Supplementary Figure 2. *In vitro* selection of SALL4-targeting sgRNA and target-specificity of CRISPR-DiR. (A)** On the left agarose gel image: sgRNA efficiency is tested for ASCL1 DNA template. The “‑” denotes a non-targeting control sgRNA for ASCL1, while “sgASCL1” refers to a validated sgRNA sequence for ASCL1 targeting. On the right agarose gel image: sgRNA efficiency is tested for the SALL4 DNA template. 11 sgRNAs were tested in total, and their target sequences are listed in Supplementary Table 4. sgRNAs 5, 6, 7, and 11 demonstrated efficient cleavage of the SALL4 DNA backbone (white box) and were shortlisted for further experiments (renamed as sgSALL4_1 to sgSALL4_4). Their respective target site and strand direction in SALL4 locus is depicted. **(B)** FACS analysis of dCas9 transduced cells. Cells expressing dCas9-mCherry were identified by red fluorescence and sorted accordingly for subsequent transduction with shortlisted sgRNAs that are conjugated with the DiR. **(C)** Bisulfite sequencing data for SALL4 exon 4 after CRISPR‑DiR in SNU387. The numbers indicate each of the CpG dinucleotide in exon 4. sgSALL4_1 refers to the efficient SALL4-targeting sgRNA that demethylates the SALL4 5’ UTR ‑ exon 1 ‑ intron 1 region in Figure 2B. **(D)** SALL4 transcript levels after CRISPR-DiR in SNU387. SALL4 mRNA levels were monitored for up to 21 days after the transduction of individual sgRNA-DiR. NT refers to sgRNA-DiR that does not contain a SALL4-targeting sequence. Mean ± SD, n ≥ 3, *P < 0.05; **P < 0.01; ***P < 0.001.

**Supplementary Figure 3. Characterization of HepAD38B, a hepatitis B-inducible HCC cell line. (A)** Absolute quantification of hepatitis B viral transcripts during hepatitis B induction in HepAD38B, including four major viral transcripts: “core” for capsid protein, “surface” for small, middle, and large surface proteins, “polymerase” for DNA polymerase, and “HBx” for hepatitis B antigen X. **(B and C)** transcript and protein levels of SALL4, SALL4P5, SALL4P7, and HBx during hepatitis B induction in HepAD38B. HBx levels were monitored for over 20 days as a positive control. SALL4 protein expression in SNU398 wildtype was used as a positive control for SALL4 immunoblotting for the HepAD38B cell line as it lacks SALL4 expression initially. Mean ± SD, n ≥ 3.

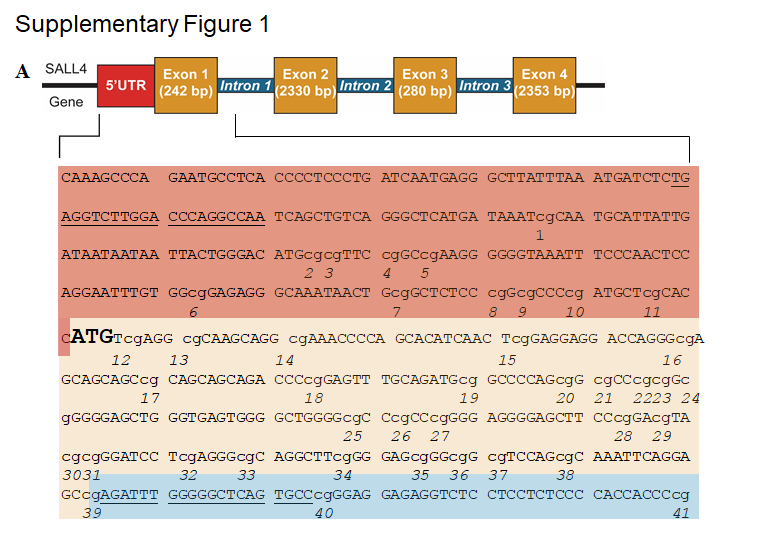

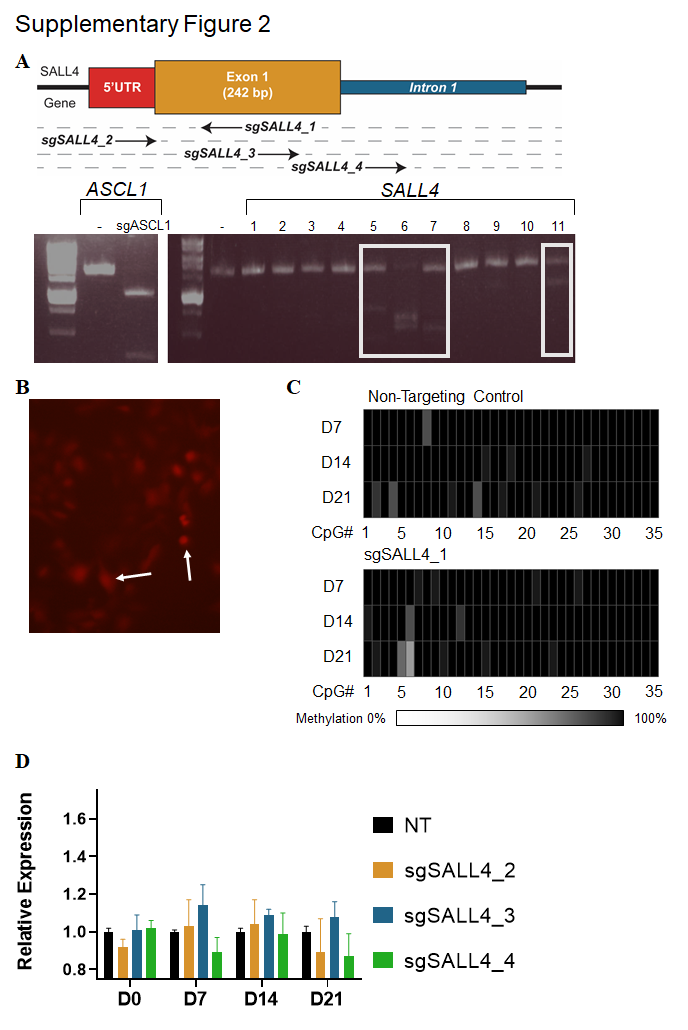

Supplementary Figure 3

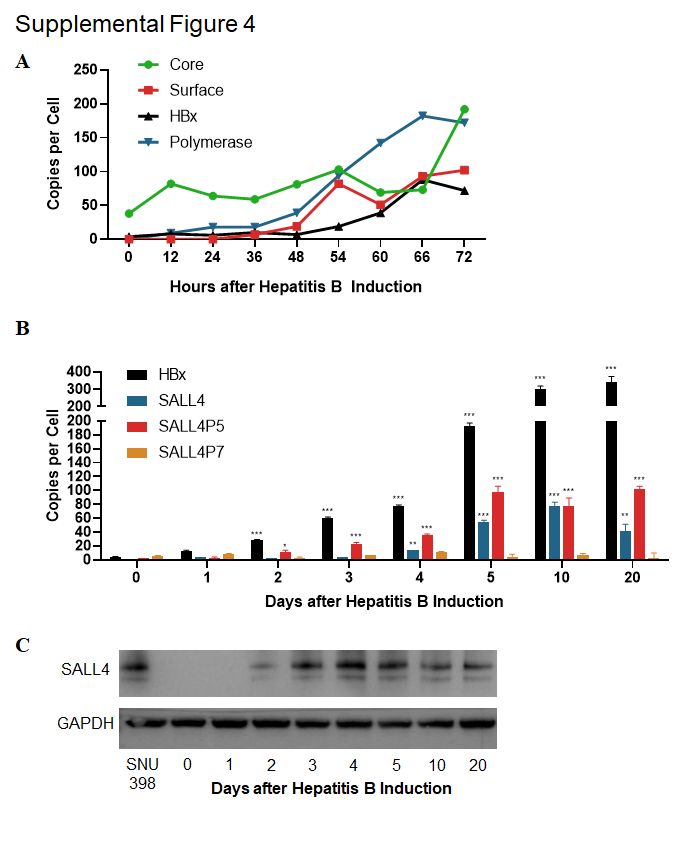

**Supplementary Tables**

**Table S1. Primer sequences used for qRT-PCR and ddPCR.**

| **Target genes** | **Direction** | **Primer sequences (5’ to 3’)** |
| --- | --- | --- |
| SALL4A | Forward | TGCAGCAGTTGGTGGAGAAC |
|  | Reverse | TCGGTGGCAAATGAGACATTC |
| SALL4B | Forward | AAGCACAAGTGTCGGAGCAG |
|  | Reverse | TCCCACAAATGTGCCAGGAA |
| SALL4P 5 | Forward | ACCACCAAAGGCAGCTTGAA |
|  | Reverse | CTGTACCTAATGGAGCCCCC |
| SALL4P7 | Forward | CACCCACACCCCAGAACATA |
|  | Reverse | TTGCCCCATAGTGCCTGAAG |
| GAPDH | Forward | GACAGTCAGCCGCATCTTCT |
|  | Reverse | GCGCCCAATACGACCAAATC |
| β-actin | Forward | GGGAGATACCATGATCACGAAGGT |
|  | Reverse | CCACAAATTATGCAGTCGAGTTTCCC |

**Table S2. Primer sequences for bisulfite sequencing**

| **Target genes** | **Direction** | **Primer sequences (5’ to 3’)** |
| --- | --- | --- |
| 5’UTR-Exon1-Intron 1 | Forward | TGAGGTTTTGGATTTAGGTTAA |
|  | Reverse | AACACTAAACCCCCAAATCT |
| Exon 4 | Forward | GGGAGGAGGGTTTAATGTTTT |
|  | Reverse | AACTCCCCTCRATCTAAAAAAC |

**Table S3. Primer sequences with the T7-tag used for PCR amplification**

| **Oligo Names** | **Direction** | **Oligonucleotide Sequences** |
| --- | --- | --- |
| SALL4  P5 sense strand | Forward | CGTTAATACGACTCACTATAGGGGAGTTAGGCTGTATGAGGA |
|  | Reverse | ACCTCAGGCAAGTCCTGCAAT |
| SALL4  P5 antisense strand | Forward | GGAGTTAGGCTGTATGAGGA |
|  | Reverse | CGTTAATACGACTCACTATAGGACCTCAGGCAAGTCCTGCAAT |

**Table S4. Primer sequences used for SALL4 and SALL4 pseudogene cloning**

| **Primer Names** | **Primer sequences** | **Restriction sites** | **Plasmid vectors** |
| --- | --- | --- | --- |
| SALL4 F | AAGCTTACATCTCCGCGGTGGATGT | HindIII | pCMV6 |
| SALL4 R | GGATCCTGCTCCGACACTTGTGCTTG | BamHI |  |
| SALL4P5 F | AAGCTTGCACCATGTTGAGGTGCAAG | HindIII | pCMV6 |
| SALL4P5 R | GGATCCTGATGCTTGTCGAGGACTGC | BamHI |  |
| SALL4P7 F | AAGCTTCCTGGACTCTGTGCTCTTCC | HindIII | pCMV6 |
| SALL4P7 R | GGATCCTGTTCCATGGCTTCCTTTTC | BamHI |  |
